# High Real-time Reporting of Domestic and Wild Animal Diseases Following Rollout of Mobile Phone Reporting System in Kenya

**DOI:** 10.1101/2020.12.04.411348

**Authors:** Kariuki Njenga, Naomi Kemunto, Samuel Kahariri, Lindsey Holmstrom, Harry Oyas, Keith Biggers, Austin Riddle, John Gachohi, Mathew Muturi, Athman Mwatondo, Francis Gakuya, Isaac Lekolool, Rinah Sitawa, Michael Apamaku, Eric Osoro, Marc-Alain Widdowson, Peninah Munyua

## Abstract

**Background:** To improve early detection of emerging infectious diseases in sub-Saharan Africa (SSA), many of them zoonotic, numerous electronic animal disease-reporting systems have been piloted but not implemented because of cost, lack of user friendliness, and data insecurity. In Kenya, we developed and rolled out an open-source mobile phone-based domestic and wild animal disease reporting system and collected data over two years to demonstrate its robustness and ability to track disease trends.

**Methods:** The Kenya Animal Biosurveillance System (KABS) application was built on the Java^®^ platform, freely downloadable for android compatible mobile phones, and supported by web-based account management, form editing and data monitoring. The application was integrated into the surveillance systems of Kenya’s domestic and wild animal sectors by adopting their existing data collection tools, and targeting disease syndromes prioritized by national, regional and international animal and human health agencies. Smartphone-owning government and private domestic and wild animal health officers were recruited and trained on the application, and reports received and analyzed by Kenya Directorate of Veterinary Services. The KABS application performed automatic basic analyses (frequencies, spatial distribution), which were immediately relayed to reporting officers as feedback.

**Results:** Over 95% of trained domestic animal officers downloaded the application, and >72% of them started reporting using the application within three months. Introduction of the application resulted in 2- to 10-fold increase in number of disease reports when compared the previous year (p<0.05), and reports were more spatially distributed. Among domestic animals, food animals (cattle, sheep, goats, camels, and chicken) accounted for >90% of the reports, with respiratory, gastrointestinal and skin diseases constituting >85% of the reports. Herbivore wildlife (zebra, buffalo, elephant, giraffe, antelopes) accounted for >60% of the wildlife disease reports, followed by carnivores (lions, cheetah, hyenas, jackals, and wild dogs). Deaths, traumatic injuries, and skin diseases were most reported in wildlife.

**Conclusions:** This open-source system was user friendly and secure, ideal for rolling out in other countries in SSA to improve disease reporting and enhance preparedness for epidemics of zoonotic diseases.

**Authors Summary:** Taking advantage of a recently developed freely downloadable disease reporting application in the United States, we customized it for android smartphones to collect and submit domestic and wild animal disease data in real-time in Kenya. To enhance user friendliness, the Kenya Animal Biosurveillance System (KABS) was installed with disease reporting tools currently used by the animal sector and tailored to collected data on transboundary animal disease important for detecting zoonotic endemic and emerging diseases. The KABS database was housed by the government of Kenya, providing important assurance on its security. The application had a feedback module that performed basics analysis to provide feedback to the end-user in real-time. Rolling out of KABS resulted in >70% of domestic and wildlife disease surveillance officers using it to report, resulting in exponential increase in frequency and spatial distributions of reports regions. Utility of the system was demonstrated by successful detected a Rift Valley fever outbreak in livestock in 2018, resulting in early response and prevention of widespread human infections. For the wildlife sector in Eastern Africa, the application provided the first disease surveillance system developed. This open-source system is ideal for rolling out in other countries in sub-Saharan Africa to improve disease reporting and enhance preparedness for epidemics of zoonotic diseases.

## Introduction

An effective animal disease surveillance system is important for early detection of endemic and emerging diseases, including zoonotic diseases that can rapidly spread to humans.[1–5] Among livestock farmers of sub-Saharan Africa (SSA), most of them inhabiting rural regions with high human and livestock interactions, early detection of livestock diseases can also reduce the socio-economic impact associated with loss of productivity or deaths of the animals.[6] Some of these rural areas are also inhabited by diverse species of wildlife, creating a conducive environment for emergence and transmission of pathogens across the wildlife-livestock-human interface.[2,7,8]

According to the Food and Agriculture Organization, effective surveillance for animal diseases in SSA is limited by absence of surveillance systems in the wildlife sector, inadequate trained human capacity in the domestic animal sector, and lack of user-friendly electronic data capture tools.[3] Whereas numerous electronic animal disease-reporting tools have been piloted in SSA, none has been roiled out because of cost, lack of user friendliness and analytical capabilities, and data insecurity.[3,9] Most of the platforms were also proprietary, increasing concerns around data security and access by countries.[3] Mobile phones offer a unique opportunity to develop data collection tools that can be easily integrated into existing surveillance systems with minimal resources.[10–13] Approximately 75% of the world’s population has access to mobile phones, including communities living in rural areas of developing countries[14,15]. In Kenya, 83.9% of the population has access to smartphone and the country leads the African continent in smartphone usage[16]. The flexibility, portability, connectivity and geo-location features of smart phones makes them an effective tool for electronic collection and submission of animal health data in real-time. Such capabilities may enhance the early detection of an emerging disease and strengthen preparedness, response, and mitigation activities.[13]

During the 2015-2016 *El Niño* rains when Kenya faced the threat of a Rift Valley fever (RVF) outbreak, the Directorate of Veterinary Service created an enhanced phonebased syndromic surveillance system targeting RVF hotspots in the country to provide early warning for the disease[17]. The system collected weekly data on RVF-like syndromes in cattle, sheep, goats, and camels from farmers located in high-risk counties during the four-month flooding season[17] We built on this initiative by developing a national mobile phone-based data collection system for Kenya, referred to as Kenya Animal Biosurveillance System (KABS), utilizing smart phone mobile technologies for animal health data collection initially developed in the United States[18]. Here, we describe how the KABS reporting application was customized and rolled out to report disease syndromes among domestic and wild animal populations in Kenya. We also present data collected during the 2017-2019 period.

## Methods

### KABS platform design

The Kenya Animal Biosurveillance system (KABS) application encompasses a mobile application for data collection, a web-based application for account management and data collection form editing, and a web-based dashboard for data viewing, monitoring, and download. We built the KABS mobile application on the Java^®^ platform for use on Android compatible mobile phones and made it available in the Google Play Store (Sun Microsystems, Santa Clara, CA). The application has four components; data collection that includes tools currently used by the Kenya livestock sector, account management, form editor, and reporting portals (**Figure 1**).

**Figure 1.**
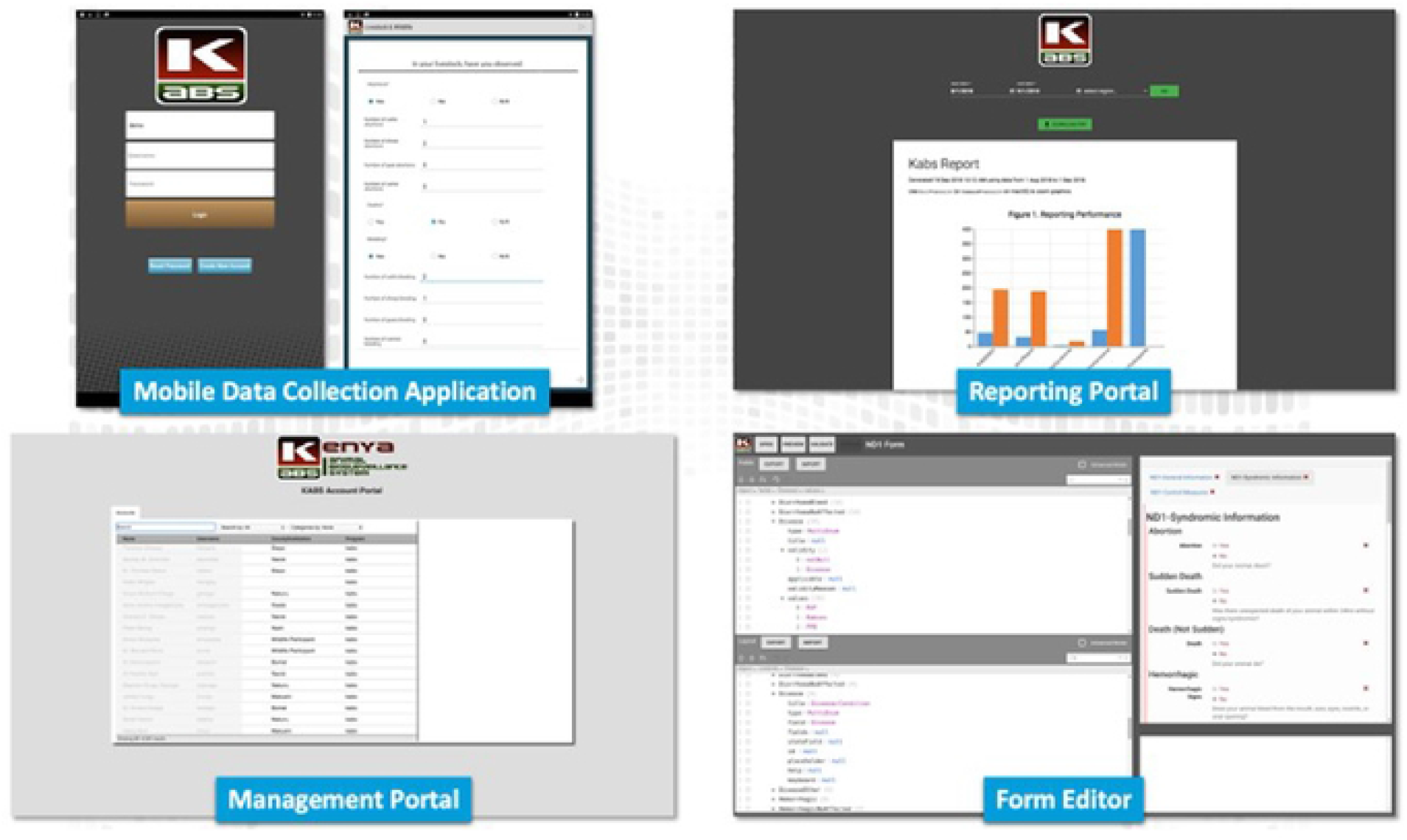
Illustration of the four components of the KABS mobile phone application platform used to report domestic and wild animal disease in Kenya.

### Customization of the application for Kenya

To integrate KABS within the disease surveillance system at the Kenya Directorate of Veterinary Services (KDVS), we added the disease surveillance forms currently used by KDVS (the ND-1 and zero report forms) in the application. Since the Kenya Wildlife Service (KWS) did not have a disease surveillance system, we jointly developed a wildlife disease-reporting tool. Working with veterinarians from the KDVS and KWS, two separate lists of domestic animals (Table 1) and wildlife (Table 2) disease syndromes for reporting were developed. Selection of reporting syndromes was guided by the World Organization for Animal Health’s (OIE) list of notifiable diseases, important transboundary animal diseases in sub-Saharan Africa, and endemic and emerging zoonotic diseases in the region.[19]

The ND-1 report form was used to report the presence of nine domestic animal disease syndromes from domestic animals (Table 1), whereas the zero report form reported the absence of important transboundary diseases including RVF, Rinderpest, Peste des petit ruminants (PPR), contagious caprine pleuropneumonia (CCPP), contagious bovine pleuropneumonia (CBPP), and avian influenza. The wildlife disease reporting form reported nine wildlife disease syndromes (Table 2). Given the diversity of wildlife species in East Africa (>2000 different animal species), we categorized them into five broad groups for reporting purposes; herbivores, carnivores, avian, aquatic mammals, and non-human primates. Under each group, important animal species were identified and included in a drop-down menu.

### Data security

The KDVS received, stored, and managed all data, with exclusive rights to grant access. Accessing the reporting portal required approval, thus limiting end-user access to data from one or more counties (i.e., for local/regional monitoring) while allowing national leaders access to nationally data. The Veterinary Epidemiology and Economic Unit at KDVS approved all requests for new accounts, which came with personal identifiers of the end-user for verification of reports. Only the senior surveillance officers of the KDVS and the Kenya Wildlife Services, and county directors of veterinary services could view all data, while sub-county officers and end-users viewed data from their localities.

### Disease data collection, transmission, and feedback to end-user

Using the KABS data collection interface, a surveillance officer could select and complete the appropriate disease report tool during any farm or wildlife conservation visit anywhere in the country. A definition of each disease syndrome was available in the surveillance tool to ensure homogeneity of data collected (Tables 1 & 2). Among data collected were geographical coordinates, animal species, number at risk and affected, and provisional diagnosis. A surveillance officer was able to enter data in KABS when in areas with no internet connectivity, allowing these data to be transmitted to the server once internet connectivity was regained.

Submitted data were received and stored at the KDVS central server at Kabete, Nairobi, and backed up on the Kenya Ministry of Agriculture, Livestock, and Fisheries central database in Nairobi. The KAB dashboard had an automated analytical tool designed to aggregate data and create spatial maps of the syndromic data, displayed as frequency histograms, tables, and maps, and automatically distributed to end-users for action, and to senior regional and national animal health officers (**Figure 1**). Due to initial bandwidth limitation, roll out of the feedback module started in January 2020. The reporting portal also provided capabilities for user-driven filtration of data to a certain date or region of the country, including downloading of the raw data for further analysis.

### User training

To be recruited and trained, animal surveillances officers from both government and private sectors needed to have a personal android-based smart phone. The five-day training included hands-on sessions to download the application, create user accounts, and input data using simulated scenarios. The KABS application was initially piloted in three counties, and feedback used to make improvements before rollout to the rest of the country. Immediately after each training session, a WhatsApp group of the users was maintained for 3 months to promote timely troubleshooting and experience sharing by system users (WhatsApp Ince, Mountain View California, USA). By the end of December 2020, we had trained domestic animal surveillance officers from 35 of the 47 counties in Kenya, and wildlife officers from all wildlife conservations in the country.

### Data analysis

We analyzed the uptake of the KABS application among the trained surveillance officers to determine it acceptance. To determine improvement in reporting domestic animal diseases, we compared reporting one year before (data collected between 2016 and 2017) and one year after (data collected between 2018 and 2019) the introduction of KABS in six counties. These data were exported into the R statistical package and compared using paired t-test.[20] In addition, descriptive analysis of domestic and wild animal disease data collected through the application was performed. No systematic wildlife disease data were collected prior to introduction of KABS to enable comparison of reporting rates.

### Ethical approval

The development and rolling out of KABS application were led by the KDVS and approved by the Kenya Ministry of Agriculture, Livestock and Fisheries as non-research.

## Results

### Uptake of KABS application

By December 2020, 697 domestic animal health officers from 35 counties had been trained on KABS, including 502 (72%) government and 195 (28%) private animal health professionals. Of those trained, 95% (N = 662) downloaded and installed the application into their smartphones. Among those who installed the application, 72% (477/662) submitted a report through the application within 3 months, including 76% government and 63% private animal health professionals. We trained 47 wildlife officers including veterinarians and research scientists; 42 (89.4%) of them from the government and five (10.6%) from private sector. All trained wildlife officers installed the application into their smartphones; and 57.4% (27/47) submitted a report through the application within 3 months.

### Spatial and temporal trends on disease reporting

To assess the impact of KABS rollout on disease reporting in domestic animals, we compared number and spatial distribution of reports received from six counties where KABS reporting had occurred for at least 12 months, including Bomet, Kilifi, Kwale, Nandi, Makueni, and Siaya. As shown in **Figure 2**, there was 2- to 10-fold increase in number of reports upon introduction of KABS across the six counties when compared to the previous year (t = −6.1053, p<0.05). Similarly, reports were more spatially distributed indicating that areas under effective surveillance had been broadened (**Figure 3**).

**Figure 2.**
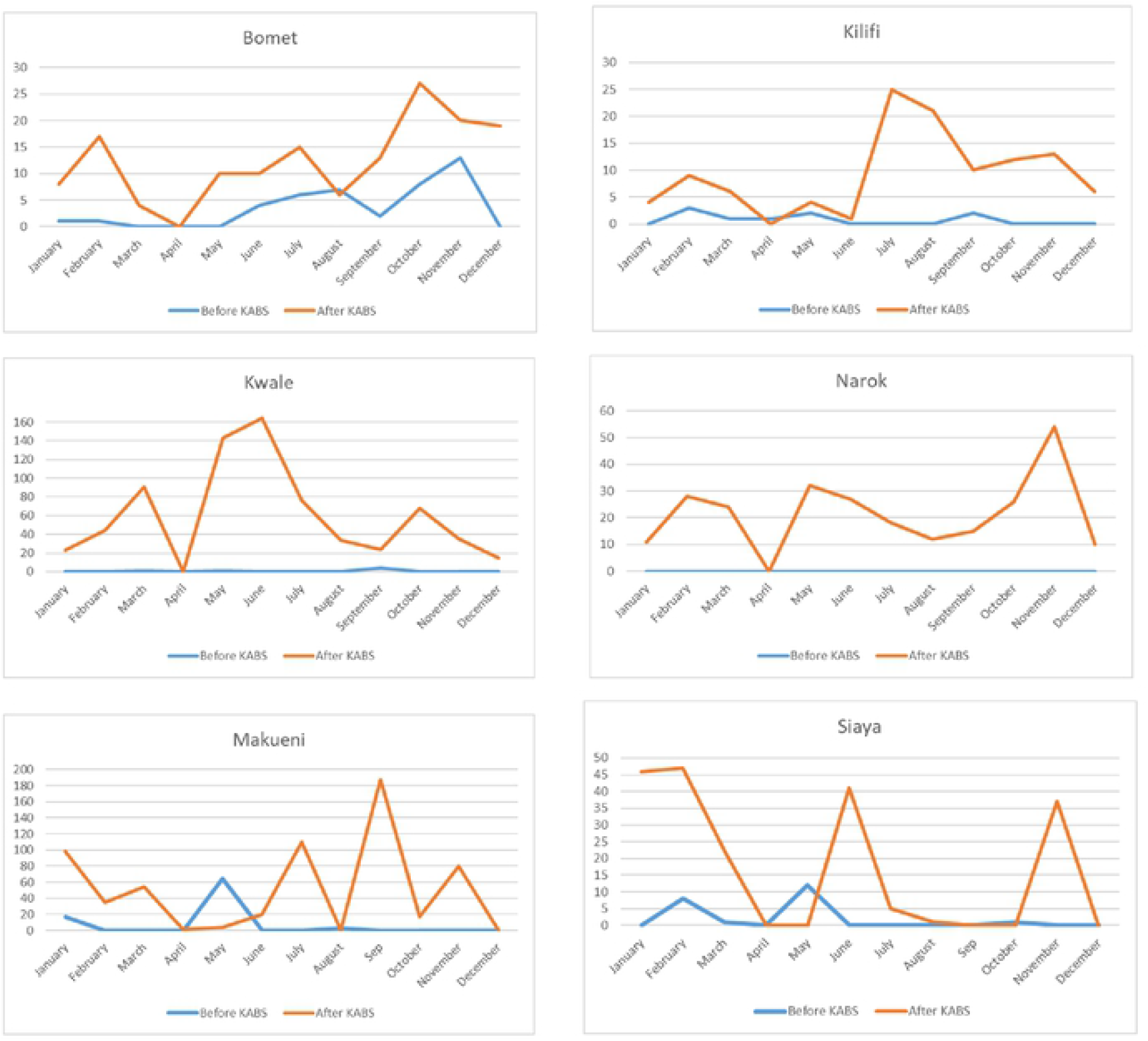
Frequency of disease reporting before (blue line) and after (red line) KABS mobile phone application introduction in Bomet, Kilifi, Kwale, Makueni, Narok, and Siaya counties of Kenya

**Figure 3.**
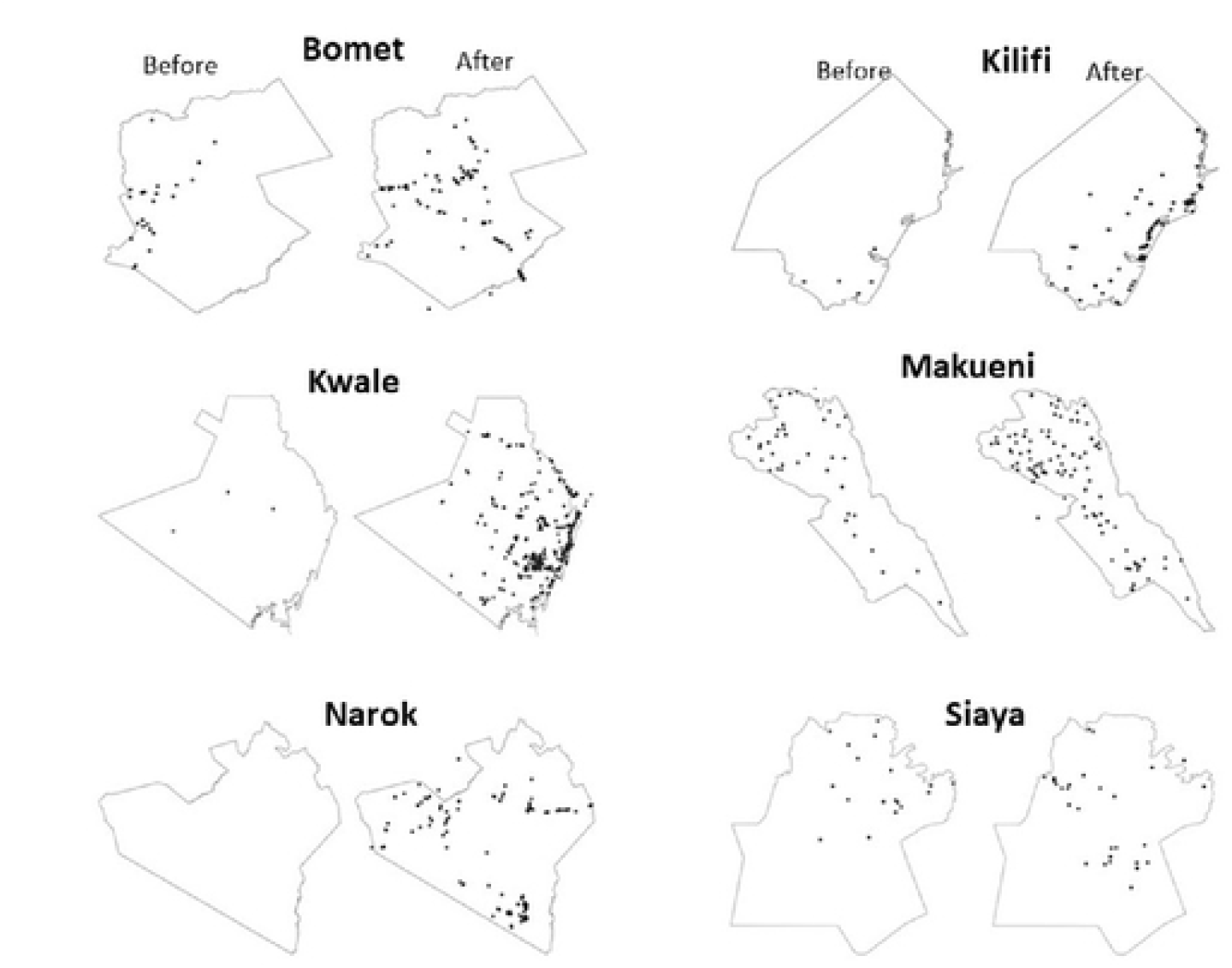
Spatial distribution of disease reporting before and after KABS mobile phone application introduction in Bomet, Kilifi, Kwale, Makueni, Narok, and Siaya Counties of Kenya.

### Summary of reported domestic and wild animal disease syndromes

As of December 2019, 11,399 domestic and 205 wild animal disease reports were received through KABS. All reports were submitted and received within 24 hours of surveillance officer collection. Livestock (cattle, sheep, goats, camels, and chicken) accounted for >90% of the reports, including cattle (51.4%), goats (25.6%), sheep (8.7%), and chicken (6.5%) (Table 3). Given that the country has >4 million camels, the number of camel disease reports was low (1%). The low number of reports among non-food animals (dogs, donkeys, and cats) and pigs also reflected their low population number in the country. In cattle, sheep, goats, and chicken, respiratory syndromes reports accounted for >50% of the reports, followed by gastrointestinal and skin conditions, and death (**Figure 4**).

**Figure 4.**
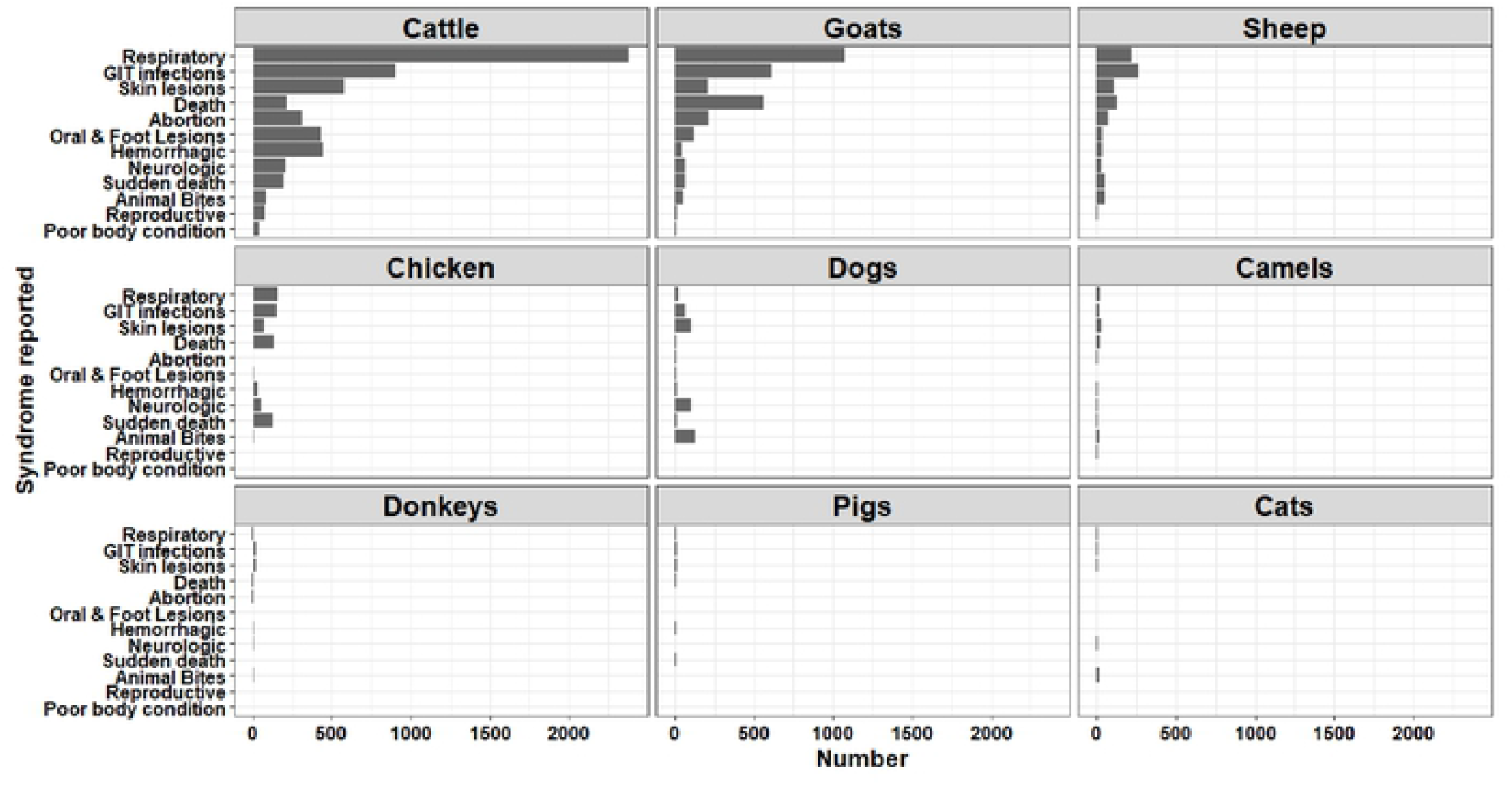
Trends in disease syndromes reported in domestic animals in Kenya using the KABS mobile phone application, (2017 – 2019).

Among wildlife, herbivores including zebra, buffalos, elephants, giraffes, and antelopes accounted for >60% of the reports, followed by carnivores (lions, cheetahs, hyenas, jackals, and wild dogs) with approximately 20% of the reports (Table 3). Deaths, traumatic injuries, and skin conditions were the most reported syndromes in wildlife (**Figure 5**).

**Figure 5.**
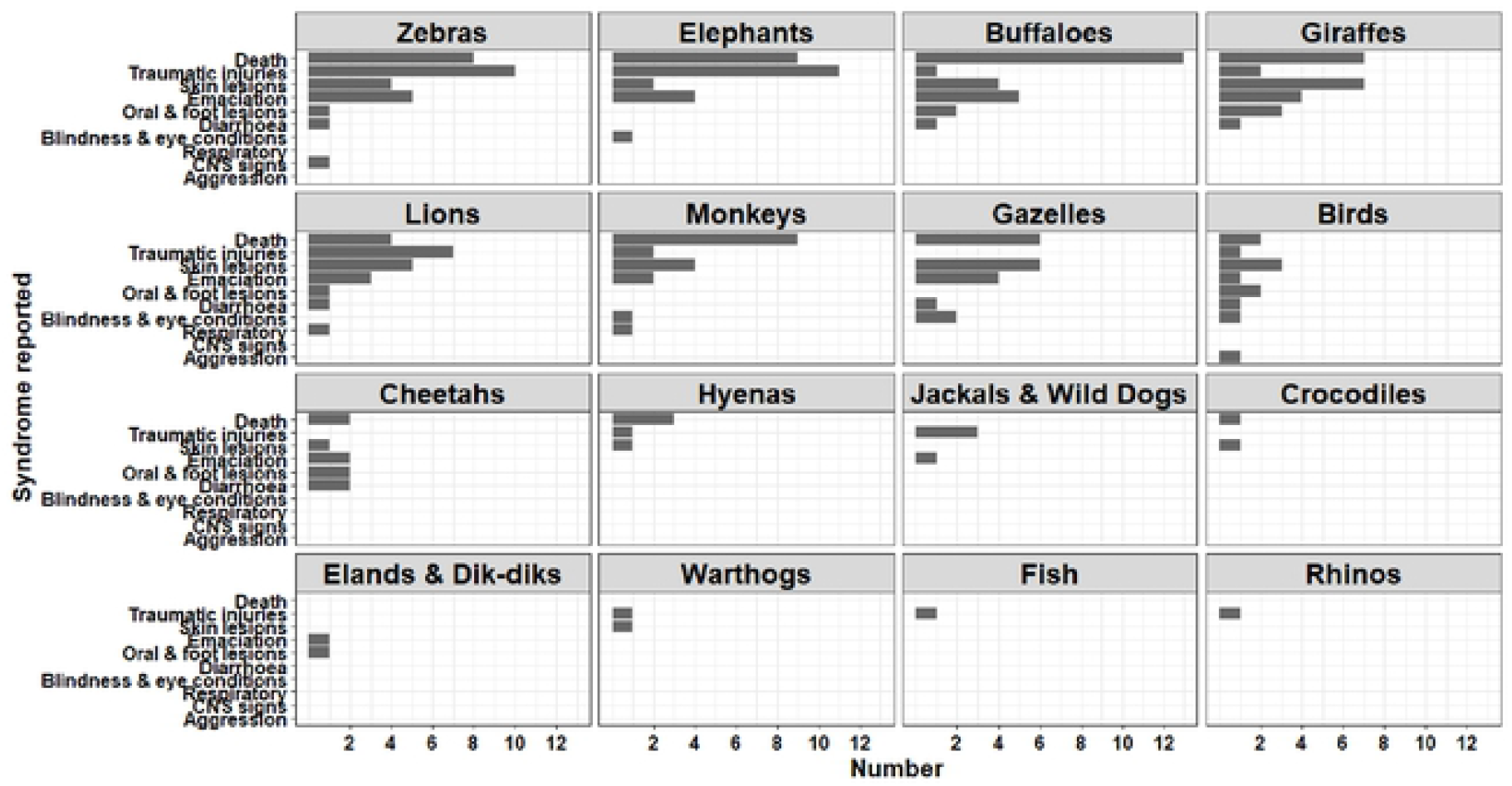
Trends in disease syndromes reported in wildlife in Kenya using the KABS mobile phone application, (2017 – 2019).

## Discussion

KABS is among the first electronic disease surveillance reporting applications to be rolled-out nationwide in SSA, providing near-real-time (within 24 hours) animal disease data collection and transmission that enhanced early detection of disease events. Within three years of introduction (2017-2020), the application had been rolled out in >70% of the country, including the entire wildlife sector. The effectiveness of application was demonstrated in 2018 when KABS detected a RVF outbreak that affected livestock and humans across three counties.[21] Disease surveillance in the domestic animal sectors in Kenya and SSA has been primarily paper-based, whereas the wildlife sectors rarely conducted routine disease surveillance and reporting. Therefore, the roll out of KABS in the wildlife sector represented a major milestone and an opportunity for detecting and responding to diseases before spillover to livestock and humans.

The high installation rate of KABS by domestic animal health officers (>95%) reflected both the acceptance of the application and dominance of android smartphones in the market. Over 70% of the officers submitted a report within three months of training, rates comparable to those observed when a similar system was rolled out in the United States.[18] Similarly in the wildlife sector, there was 57% reporting by veterinarians and research officers. By using their own phone for surveillance, and data capture tools routinely used, the application became widely accepted by all cadres of animal health professionals interested in reporting disease. The rapid submission of data and automated analytic capability allowed generation of reports for immediate feedback to the end users for action. This feedback was a motivation to keep reporting. [18]

For the government of Kenya, KABS addressed several long-standing challenges. First, the application was open-source and free, removing the cost constraints presented by previous applications. The only cost was associated with customizing the application to the Kenya animal disease surveillance system and training. Second, the data were stored and managed by the government of Kenya, providing important assurance to the country on ownership and security. Third, the application was user friendly, in part because the data collection tools were familiar, and syndromes selected for reporting covered important and notifiable transboundary animal diseases that the government was required to report. Reports were submitted immediately, eliminating the lag time experienced with other systems that require an additional step of data compilation, and thus enhancing Kenya’s ability to submit data to the OIE in a timely manner.[3,22]^-24^ Further, the application worked well in remote areas with no internet connectivity, allowing login and data collection while offline and automatic transmission of results to the KDVS once connectivity was restored. The automated collection of geographic coordinates of reported disease events enhanced follow-up by the county and national animal health officers.

The rollout of KABS resulted in up to 10-fold increase in reporting and an exponential increase in spatial distribution of reports. In contrast with the previous animal disease reporting system that relied on government officers, we recruited both government and private animal health professionals as surveillance officers, resulting in an increased rate of reporting and a broader spatial distribution of reports. In fact, we think that the increase was limited by the number of surveillance officers trained in each county. Our team is developing a training-of-trainers program that provides training materials to the current surveillance officers in each county to encourage them to train others for broader usage. The cumulative monthly reporting across the country varied, increasing during the April to June and September to November rainy seasons, and decreasing during the December to March and July to August dry seasons (Figure 2). This trend is similar to those reported over the years by the government of Kenya; however, the overall magnitude of reports has increased.[23] The low reporting in certain areas (e.g. western Kilifi) was low because the region is not inhabited by humans and livestock but instead occupied by a national park.

As shown in Figure 4 the number of reports were higher for food animals, particularly cattle, goats, sheep, and chicken than for non-food animals (dogs, cats, donkeys). Similar trends have been reported in other studies, associated with farmers attaching greater social-economic value to these animals.[24] Among wildlife, the variable number of reports across species is likely due to several factors including the size of animal populations, higher number of surveillance officers in some conservation areas, and the ease of sighting and observing some species. Thus, it was easier to sight, observe and report disease conditions in larger terrestrial wild mammals (elephants, buffalos, zebras, giraffes, lions) than in birds and aquatic animals. Respiratory, gastrointestinal, and cutaneous syndromes were the most reported among all domestic animals, whereas death, cutaneous lesions, traumatic injuries and emaciations were the most common conditions among wildlife.

The project had some limitations. Adding a testing and diagnosis module to the application will improve its utility in both domestic and wild animal sectors. Broader use of the application in SSA can be enhanced by linking KABS reports to the human health system through the emergency operation centers, thus providing early warning of potential outbreaks of zoonotic infectious diseases from both domestic and wild animals. To realize this step, standardization of syndromic and disease classification across countries and between the human and animal sectors would need to be achieved.[25] The low human capital in the wildlife sector was a setback for increasing the number of disease reports from that sector.

## Authors contribution

KN, NK, SK, PM, and LH conceived, designed, and coordinated the project. This team also conducted data analysis and developed the first draft of the manuscript. HO, JG, EO, LH, KB, and AR were involved customization and upgrading of the application. FG, IL, MM, and AM were involved in the rolling out the application including trainings and data collection. RS, MA, and MAW reviewed the data, and were involved in revising and finalizing the manuscript.

## Acknowledgements

We thank the Kenya government domestic and wild animal health officers for supporting the development and rolling out of application. The willingness of the senior leadership at the Kenya Directorate of Veterinary Services to adopt the KABS application as the primary disease-reporting tool was critical. The project was funded by US Center for Disease Control and Prevention (CDC), Grant # GH0001717 under the Global Health Security Agenda, and Food and Agriculture Organization (FAO)-Kenya Office. The CDC and FAO scientists listed as coauthors were involved in the design and rolling out of KABS application, including data interpretation, and manuscript writing. However, the funder had no role in the data collection and analysis for this manuscript.

## Disclaimer

The findings and conclusions in this report are those of the authors and do not necessarily represent the official position of the United States’ Centers for Disease Control and Prevention, United States’ Department of Agriculture or the Government of Kenya.

